# Sense of absence: Spatial perception through active sensing by insect antennal mechanosensory system

**DOI:** 10.1101/2021.02.23.432611

**Authors:** Nwuneke Okereke Ifere, Hisashi Shidara, Nodoka Sato, Hiroto Ogawa

## Abstract

Animals perceive their surroundings by using various modalities of sensory inputs to navigate their locomotion. Nocturnal insects such as crickets use mechanosensory inputs mediated by their antennae to navigate under dark conditions. Active sensing with voluntary antennal movements improves spatial information, but it remains unclear how accurately the insects can perceive the surrounding space by using their antennal system. Crickets exhibit escape behavior in response to a short air-puff, which is detected by the abdominal mechanosensory organ called cerci and is perceived as a “predator approach” signal. We placed objects of different shapes at different locations with which the cricket actively made contact using its antenna. We then examined the effects on wind-elicited escape. The crickets changed their movement trajectory depending on the shape and location of the objects so that they could avoid collision with these obstacles even when the escape behavior was triggered by another modality of stimulus. For instance, when a wall was placed in front of the crickets so that it was detected by one side of their antenna, the escape trajectory in response to a stimulus from behind was significantly biased toward the side opposite the wall. However, if the antenna on the free side without the wall was ablated, this modulation to avoid collision diminished, suggesting that the antenna on the free side provided information of “absence” of obstacles. This study demonstrated that crickets were able to perceive spatial information, including the presence or absence of objects by active sensing with their antennal system.

**Summary Statement:** Crickets can acquire spatial information such as shape, location and orientation of objects through active sensing by antennal mechanosensory system, which also provides information about the absence of objects.

## INTRODUCTION

Animals perceive their surroundings using various sensory inputs to navigate their locomotion appropriately. In situations where visual cues are not available, such as in dark conditions or in very tight spaces due to surrounding objects, mechanosensory inputs provide effective cues to guide their path. For example, rodents employ their facial whisker as a tactile sensor to guide their locomotion (Prescott et al., 2011; Grant et al., 2018). In insects, the mechanosensory cues provided by antennae play an important role in guiding their locomotion. For example, cockroaches walk along a wall by making contact with the wall using their antenna (Camhi and Johnson, 1999; Baba et al., 2010). These facts suggest that insects use mechanosensory inputs mediated by their antennae for appropriate navigation in various environments. However, it remains unclear whether the insect antennal system can actually perceive the surrounding space.

Here, the term “spatial perception” is taken to imply the ability of internal representation of the surrounding space, which would require the integration of spatial information of the surrounding objects. To confirm the ability for spatial perception, it is necessary to examine whether the oriented behavior induced by another stimulus, rather than the behavior directly induced by the detected objects, is modulated by the spatial context, such as the arrangement of objects. This is because, if the animal is reflexively oriented to the object itself that causes the action, it can change its behavior depending on the position of the object without spatial perception. For example, the impacts of antennal stimuli on phonotaxis in female crickets have been investigated. Active contact with an object by the antennae of the crickets performing phonotaxis reduces forward velocity toward the sound source and suppresses phonotactic steering depending on the side where object is located and the distance (Haberkern and Hedwig, 2016). This finding suggests that the spatial perception of the cricket antennal mechanosensory system may affect the oriented behavior elicited by other modalities of sensory inputs. To address this issue, we used their escape behavior in response to short airflow, which was detected by the cerci, an abdominal mechanosensory organ (Gras and Hörner, 1992; Tauber and Camhi, 1995; Oe and Ogawa, 2013).

The escape behavior is a distinctly oriented locomotion in which animals move in the opposite direction to threats, such as predators, in order to increase, as much as possible, its distance from the threat (Card and Dickinson, 2008; Dominici et al., 2011a,b). Escape trajectories are often modulated depending on the environmental context (Domenici, 2010; Evans et al., 2019). When goldfish visually perceive an obstacle, they change their escape trajectory to avoid collision with it (Eaton and Emberley, 1991). The escape directionality of lizards and mice also depends on the presence and location of shelter, detected visually and auditorily (Hennig, et al., 1976; Vale et al., 2017). The combination of mechanosensory and visual cues results in the escape of rockpool prawn over longer distances and with greater directionality when compared to those triggered by mechanosensory stimulus alone (Guerin and Neil, 2015). Also, in wind-elicited escape behavior, the crickets alter their escape trajectory elicited by a short air-puff depending on the acoustic context represented as different sound frequencies (Fukutomi et al., 2015; Fukutomi and Ogawa, 2017). In addition, the wind stimulus applied to cockroaches that make contact with the wall using their antenna elicits different turning behavior when compared to animals that do not make contact (Ritzmann et al., 1991). Thus, the modulation of the wind-elicited escape behavior by different modalities of sensory inputs would be an ideal model to test the ability of the antennal system in spatial perception.

For spatial perception, “active sensing” is one of the most reliable ways to acquire contextual cues from the environment. In mechanosensory active sensing, the animals voluntarily move their sensory organs across the surrounding objects to acquire information about their environment and to enhance the searching space and sampling frequency. Rodents use their facial whiskers not only as passive sensory organs, but also actively move them to perceive surrounding objects (Mitchinson et al., 2007; Deutsch et al., 2012; Voigts et al., 2015; Bush et al., 2016). Insects move their antennae to actively sample the surrounding space and are able to identify obstacles, recognize conspecifics and predators, actively track objects, and probe surface textures (Staudacher et al., 2005; Okada and Toh, 2006). In this study, the crickets were tethered on an air-lifted treadmill, and objects of different shapes—a cylindrical rod or a plate—was placed at different distances and in different positions with respect to the antennae. In this condition, the tethered crickets were able to detect the object by “actively” making contact with the object using their antenna. We compared the escape running triggered by an airflow stimulus between the different conditions with and without antennal stimulation. We found that the crickets could change their escape trajectory depending on the shape and position of the obstacles, and that their antennae were also necessary to perceive a free space where obstacles were absent on the escape path. This suggests that insects may be able to recognize “absence” through active sensing.

## MATERIALS AND METHODS

### Animals

We used the wild-type strain of field crickets for this study (*Gryllus bimaculatus* De Geer, 1773, Hokudai WT; Watanabe et al., 2018). The laboratory-bred adult male crickets (0.50–1.00 g body weight) were used in all the experiments. They were reared under 12 h light:12 h dark conditions at a constant temperature of 27°C. The guidelines of the Institutional Animal Care and Use Committee of the National University Corporation, Hokkaido University, Japan, specify no particular requirements for the treatment of insects in experiments. Before commencing the experiments, all crickets were checked to ensure that the legs, cerci, and antennae were intact.

### Treadmill system

We monitored a cricket’s locomotion in response to an air-puff stimulus with the same spherical-treadmill system as used in our previous studies (Oe and Ogawa, 2013; Fukutomi et al., 2015; Fukutomi and Ogawa, 2017). An animal was tethered on top of a Styrofoam ball using a pair of L-shaped insect pins that were stuck to the cricket’s tergite with paraffin wax. The cricket’s running activity was monitored as rotation of the ball at a sampling rate of 200 Hz, using two optical mice that were mounted orthogonally around the ball. TrackTaro software (Chinou Jouhou Shisutemu Inc., Kyoto, Japan) was used to measure the moving trajectory and to calculate parameters such as translational and angular turn velocities based on the measured ball rotation.

### Air-puff and tactile stimulations

An airflow stimulus was provided to the cricket that was stationary for more than one second by a short puff of nitrogen (N_2_) gas from a plastic nozzle (15 mm diameter) connected to a PV820 pneumatic picopump (World Precision Instruments, Sarasota, FL, USA). By adjusting the delivery pressure of the picopump, the velocity of the air-puffs was controlled at 0.68 m s^-1^, which was measured at the center of the treadmill with a 405-V1 thermal anemometer (Testo, Yokohama, Japan). The duration of the air-puff stimulus was set to 200 ms. Eight air-puff nozzles were arranged around the inside wall of the arena, and the height of the nozzles were aligned with the same horizontal plane as the animal. The nozzle ends were positioned at a distance of 130 mm from the center of the treadmill and were spaced 45° apart. In the experiments to test the effects of antennal tactile stimulation, the air-puff stimulus was applied from either the posterior (180°) or lateral side (90°) of the animal. In a preliminary experiment to test the effects of bilateral ablation of antennae on the escape behavior, the stimulus was provided from eight directions at an interval of 135° (Fig. S1).

A vertical cylindrical rod (ø = 7 mm) or square plate (50 mm × 50 mm) was placed in different orientations and distances from the antennae. The objects were placed either in the “far” position, which was 5 mm proximal from the tip of the antenna or in the “near” position, which was at half the antenna length (Figs 1A and 2A). For different orientations of the object, the plate was placed either on the left side or in front of the cricket in the near position. The lateral plate was placed either at the anterior or posterior position (Fig. 3A). The frontal plate was placed aligning to the center of the cricket or was placed on one side of the cricket (Figs 4A and 5A).

**Fig. 1.**
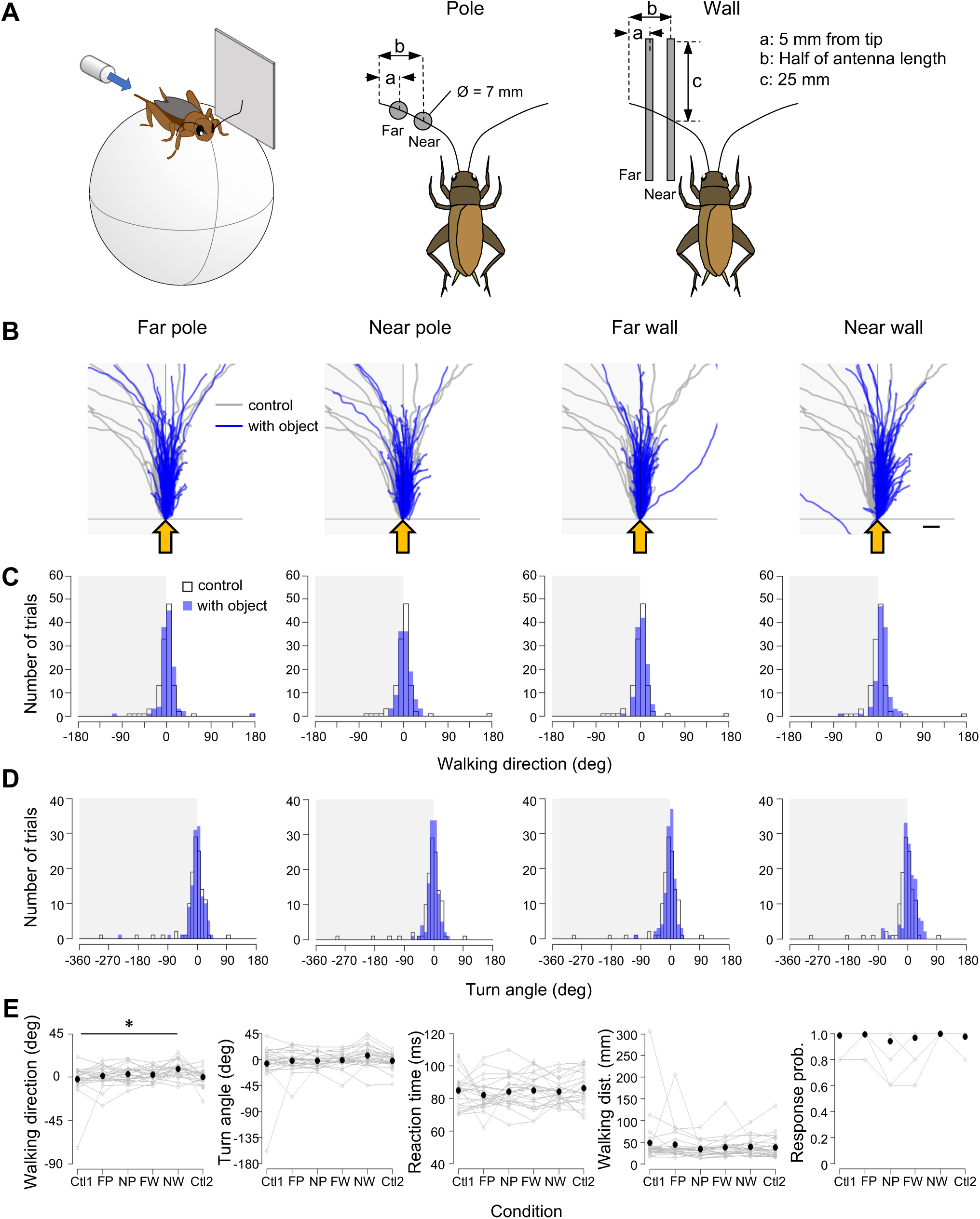
Cross-modal effects of antennal mechanosensory inputs on the escape response to airflow from behind. **A**. Experimental design. A cylindrical pole (ø = 7 mm) or square plate (50 × 50 mm) was positioned at the anterolateral position at different distances from the cricket. Far: 5 mm from the tip of the antenna. Near: Half of the antenna length. A puff of air was applied from behind the cricket. **B**. Walking trajectories in the initial response to the air puff combined with the antennal stimulation. Gray traces show the trajectories under control condition with no objects. Blue traces show the trajectories under antennal stimulation conditions. Shaded region indicates the side on which the objects were placed. Yellow arrows indicate the direction of the air puff. Scale bar indicates 10 mm. **C, D.** Distributions of walking direction (C) and turn angle (D) under different conditions. Open and blue bars indicate the data from the control and antennal-stimulation conditions, respectively. **E.** Locomotion parameters including walking direction, turn angle, reaction time, walking distance, and response probability under different conditions. N = 24 individuals. Five trials were implemented for each condition for each individual. Black dots indicate the average of the mean values for all trials for each individual. Data for the control condition were obtained for each individual twice, at the beginning (Ctl1) and at the end (Ctl2) of the experiment. FP, far pole; NP, near pole; FW, far wall; NW, near wall. * *P* < 0.05 (Fisher’s nonparametric test or Wilcoxon signed-rank test with Bonferroni correction).

**Fig. 2.**
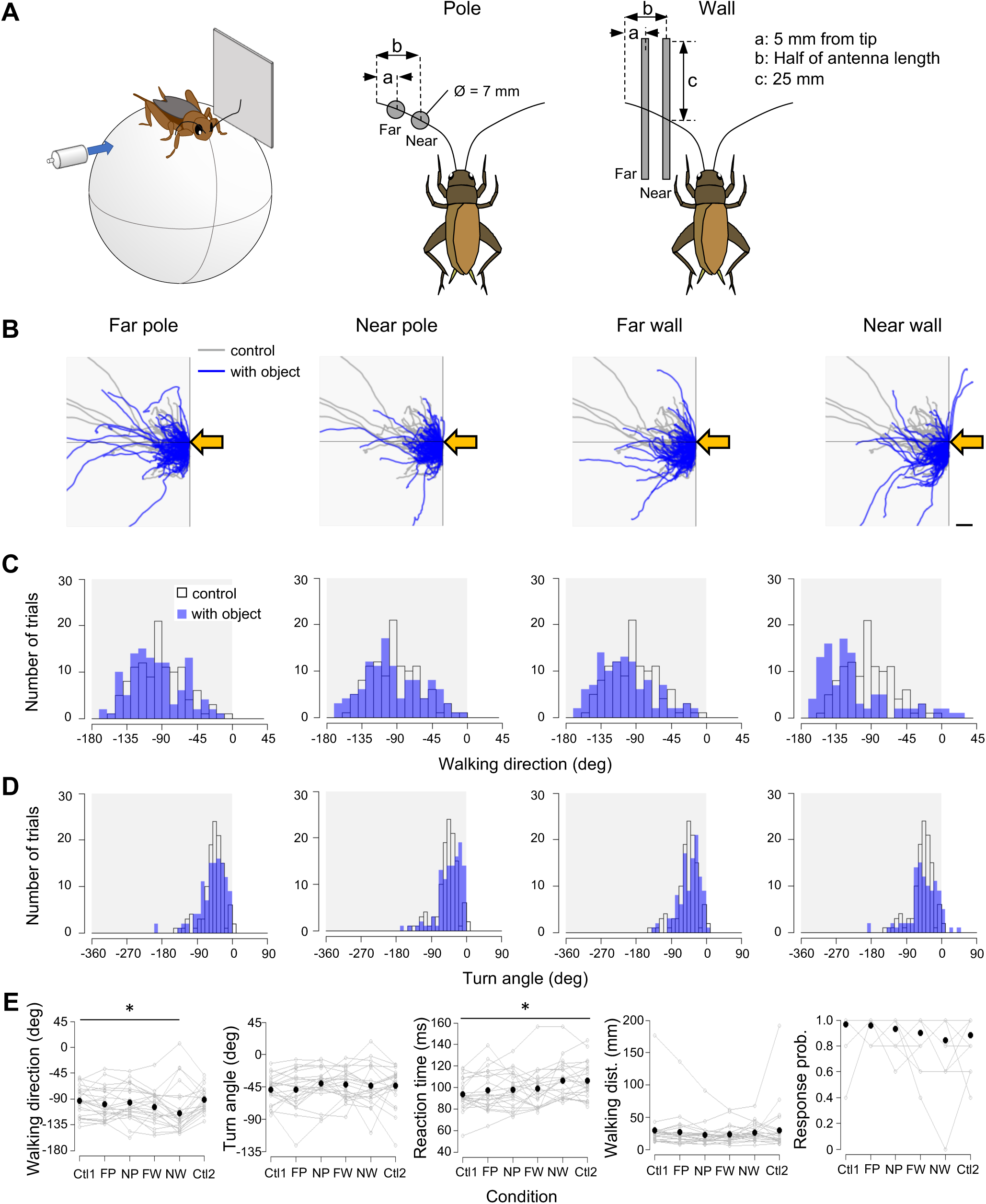
Cross-modal effects of antennal mechanosensory inputs on the escape response to airflow from the side contralateral to obstacles. **A**. Experimental design. As in figure1, a cylindrical pole (ø = 7 mm) or square plate (50 × 50 mm) was positioned at the anterolateral position at different distances from the cricket. Far: 5 mm from the tip of the antenna. Near: Half of the antenna length. Air puff was applied from the side contralateral to the objects. **B**. Walking trajectories in the initial response to the air puff combined with the antennal stimulation. Gray and blue traces show the trajectories under control and antennal-stimulation conditions, respectively. Shaded region indicates the side on which the objects were placed. Yellow arrows indicate the direction of the air puff. Scale bar indicates 10 mm. **C, D.** Distributions of walking direction (C) and turn angle (D) under different conditions. Open and blue bars indicate the data from the control and antennal-stimulation conditions, respectively. **E.** Locomotion parameters under different conditions. N = 24 individuals. Five trials were implemented for each condition for each individual. Black dots indicate the average of the mean values for all trials for each individual. Data for the control condition were obtained for each individual twice, at the beginning (Ctl1) and at the end (Ctl2) of the experiment. FP, far pole; NP, near pole; FW, far wall; NW, near wall. * *P* < 0.05 (Fisher’s nonparametric test or Wilcoxon rank-sum test with Bonferroni correction).

**Fig. 3.**
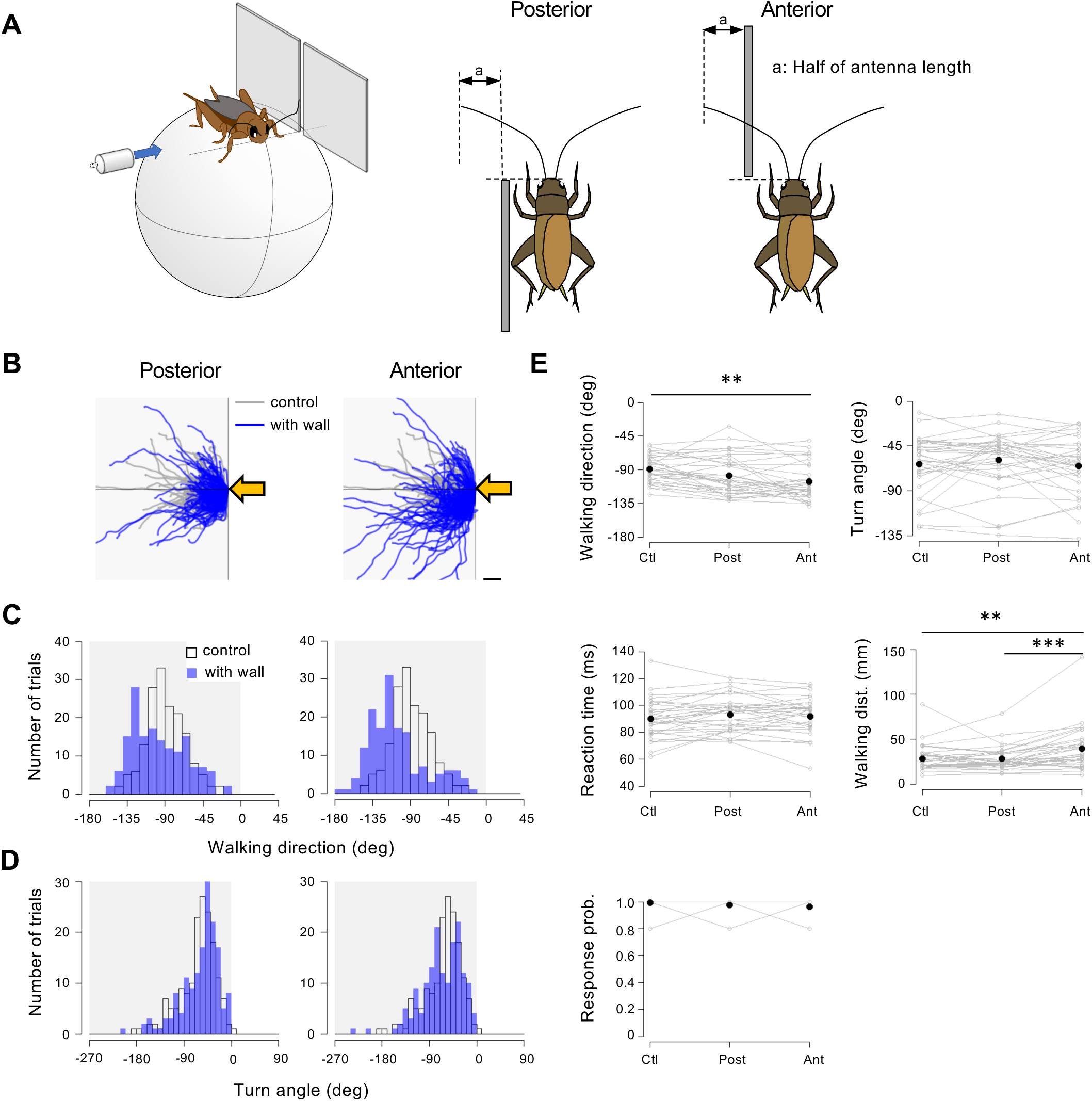
Positional effects of the wall placed laterally on the wind-elicited escape behavior. **A**. Experimental design. A square plate (50 × 50 mm) was positioned laterally at the anterior or posterior side of the cricket. A puff of air was applied from the side contralateral to the plate. **B**. Walking trajectories in the initial response to the air puff combined with the antennal stimulation. Gray and blue traces show the trajectories under the control and antennal-stimulation conditions, respectively. Shaded region indicates the side on which the plate was placed. Yellow arrows indicate the direction of the air puff. Scale bar indicates 10 mm. **C, D.** Distributions of the walking direction (C) and turn angle (D) under different conditions. Open and blue bars indicate data from the control and stimulation conditions, respectively. **E.** Locomotion parameters under different conditions. N = 33 individuals. Five trials were implemented for each condition for each individual. Black dots indicate the average of the mean values for all trials for each individual. Ctl, control; Post, posterior; Ant, anterior. ** *P* < 0.01, *** *P* < 0.001 (Fisher’s nonparametric test or Wilcoxon signed-rank test with Bonferroni correction).

**Fig. 4.**
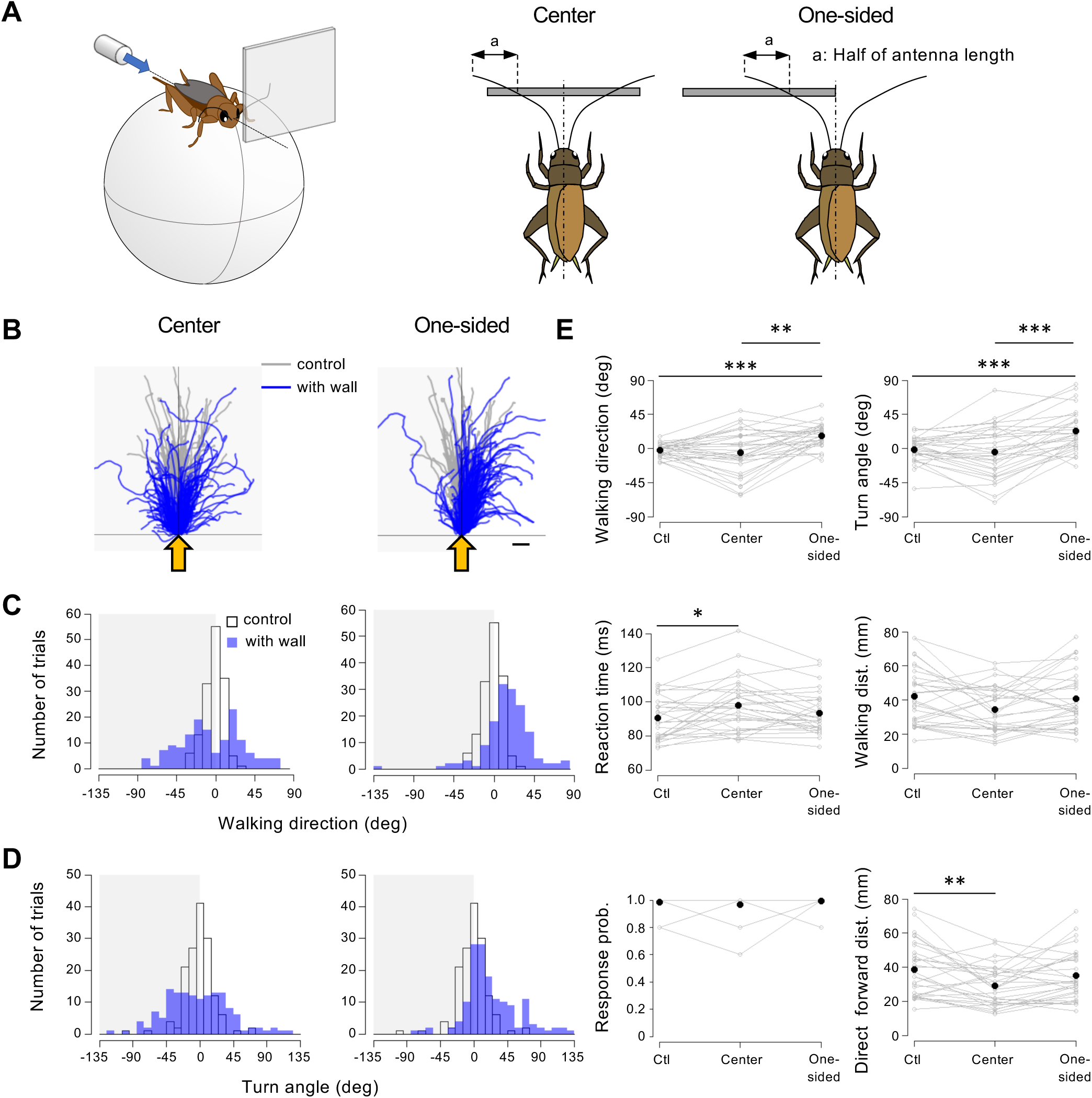
Positional effects of the wall on the wind-elicited escape behavior when the wall was placed in front of the cricket. **A**. Experimental design. A square plate (50 × 50 mm) was positioned in front of the cricket either at the center or toward one side. A puff of air was applied from behind the animal. **B**. Walking trajectories in the initial response to the air puff combined with the antennal stimulation. Gray and blue traces show the trajectories under the control and antennal-stimulation conditions, respectively. Shaded region indicates the side on which the plate was placed. Yellow arrows indicate the direction of the air puff. Scale bar indicates 10 mm. **C, D.** Distributions of walking direction (C) and turn angle (D) under different conditions. Open and blue bars indicate data from the control and stimulation conditions, respectively. **E.** Locomotion parameters under different conditions. N = 30 individuals. Five trials were implemented for each condition for each individual. Black dots indicate the average of the mean values for all trials for each individual. Ctl, control. * *P* < 0.05, ** *P* < 0.01, *** *P* < 0.001 (Fisher’s nonparametric test and Wilcoxon signed-rank test with Bonferroni correction).

**Fig. 5.**
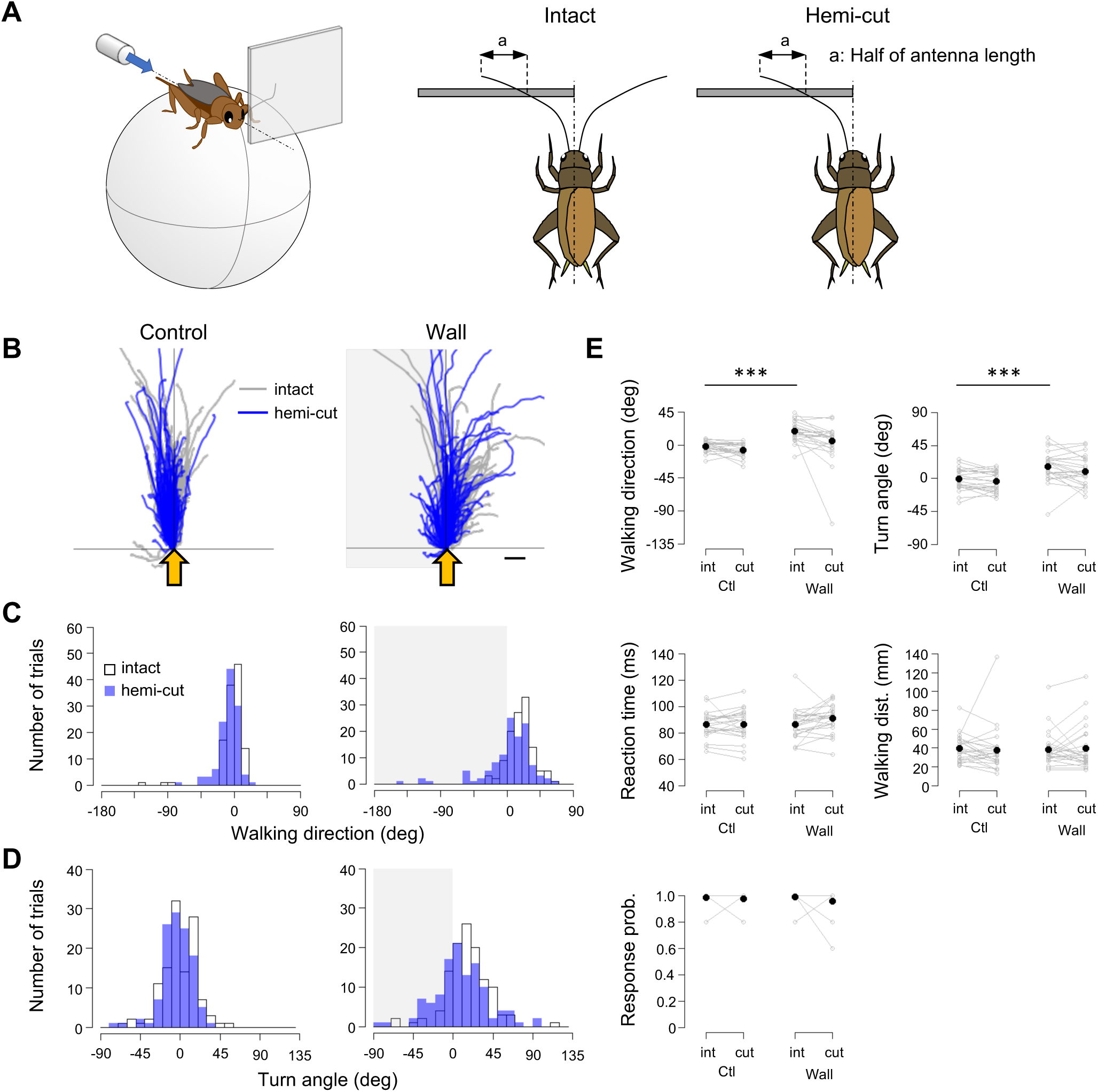
Hemi-lesion of the antenna diminished the modulation of escape trajectory when an obstacle was placed in front of the animal. **A**. Experimental design. A square plate (50 × 50 mm) was positioned on one side in front of the cricket. A puff of air was applied from behind the animal. In hemi-cut condition, the antenna contralateral to the plate was resected. **B**. Walking trajectories in the initial response to the air puff combined with the antennal stimulation. Gray traces show the walking trajectory of intact crickets. Blue traces show the walking trajectory of antenna-ablated crickets. Shaded region indicates the side on which the plate was unilaterally placed. Yellow arrows indicate the direction of the air puff. Scale bar indicates 10 mm. **C, D.** Distributions of the walking direction (C) and turn angle (D) under different conditions. Open and blue bars indicate data in intact and antenna-ablated crickets. **E.** Locomotion parameters under different conditions. N = 24 individuals. Five trials were implemented for each condition for each individual. The black dots indicate the average of the mean values for all trials for each condition. Ctl, control; int, intact. *** *P* < 0.001, (Fisher’s nonparametric test and Wilcoxon signed-rank with Bonferroni correction).

### Video recording of antennal movement

To check the frequency and duration of contact of the antenna with the object, the antennal movement was monitored using a high-speed digital camera (CHU30-B, Shodensha, Osaka, Japan) under red LED illumination. The voluntary movement of the antenna was recorded for 60 s in each trial of measurement at a frame rate of 90 frames s^-1^ with a resolution of 640 × 480 pixels. We manually counted the frames in which the antenna was in contact with the plate, and calculated the frequency and duration of contact. We compared the antennal contacts to the anterior and posterior plates positioned on the lateral side of the animal, and also the contacts of ipsi- and contra-lateral antennae to the plate positioned on one side in front of the animal. For each condition, five trials of the measurement for 1 min were performed with an interval of 1 min between trials. Five individuals were recorded in total for each experiment.

### Experimental procedure

To examine the contribution of the antennal inputs induced directly by airflow stimulus to the wind-elicited behavior, we recorded the movement of crickets with intact antennae and those in which the antenna was bilaterally ablated at the base (Fig. S1). Nothing was placed around the crickets to prevent tactile stimulation of their antennae. In this experiment, a single air-puff was delivered from each nozzle positioned at 0°, 135°, –90°, 45°, 180°, –45°, 90°, and –135°, in this order, with an inter-trial interval greater than one minute. For each individual, 40 trials (five trials for each stimulus angle) were recorded before and after the ablation of the antenna, respectively. To ensure that the animal had recovered from the damage of the antennal ablation, the experiments using the antenna-ablated crickets were performed more than 40 min after the ablation. Twenty-six individuals were tested in total.

To test the effects of object shape and distance, we adopted the following five types of stimulation conditions (Figs 1 and 2). The rod was placed near or far from the antenna, referred as “near pole” and “far pole”. The plate was placed near or far from the antenna on the side of the cricket, referred as “near wall” and “far wall”. In the control, neither rod nor plate was placed (control). The air puff was applied from the rear of the cricket or from its lateral side opposite to the objects. For each tactile stimulation condition, 10 trials were performed with an inter-trial interval greater than one minute. The stimulation conditions were tested in the following order, control, far pole, near pole, far wall, and near wall. Finally, the control condition was tested again to confirm that the escape movement was not adapted through the series of the experiment. The trials in this condition were referred to as “control2”. In total, 60 trials were performed for each individual. Twenty-four individuals were tested.

To check the turbulence effect of the air-puff stimulation when the plate was positioned on the lateral side of the cricket, we measured the escape behaviors of the cricket in which both antennae were ablated from the base (Fig. S2). After more than 40 min of antennal ablation, the escape responses to the air-puff from behind (180°) or lateral side opposite to the plate (90°) were recorded in two different conditions— without wall (control) and with wall placed at the near position. For each individual, 20 trials (10 trials for each stimulus angle) were recorded. Twelve individuals were tested in total.

To test the effects of the object location, we adopted two types of experiments (Figs 3 and 4). In the first type, the plate was placed at the lateral side of the cricket either anteriorly or posteriorly relative to its head, referred as “anterior” and “posterior”, respectively, and the air-puff was applied from the contralateral side of the plate (Fig. 3). The side edge of the plate was aligned with the base of the antenna. Three stimulation conditions, control, anterior, and posterior were tested in random order for each individual. In the second type of the experiment, the plate was placed in front of the cricket either aligned to its center or placed toward one side, referred as “center” and “one-sided”, respectively, and the air puff was applied from its rear (Fig. 4). In the center stimulation condition, the center of the plate was aligned to the midline of the animal so that the cricket could access the plate with both antennae. In the “one-sided” condition, the side edge of the plate was aligned to the midline of the animal so that the cricket accessed the plate only with the antenna ipsilateral to the plate. The three stimulation conditions, control, center, and one-sided, were tested in random order for each individual. In both types of experiments, five trials were performed for each stimulation protocol with an inter-trial interval greater than one minute. In total, 15 trials were performed for each individual for each type of experiment. Thirty-three and 30 individuals were used in the first and second types of experiments, respectively. For the one-sided stimulation condition in the second type of experiment, the plate was placed on the right side of the animals in 15 crickets, and in another 15 crickets on the left side to counterbalance the stimulation to the antenna bilaterally.

To examine the contribution of bilateral antennae to the behavioral modulation, we adopted the one-sided stimulation condition mentioned above to the unilaterally-antenna-ablated crickets (Fig. 5). An air-puff stimulus was applied from the rear of the cricket. The movement of the intact crickets was monitored in the “control” condition without the plate and in the “wall” condition where the plate was placed on one side in front of the cricket. Then, one side of the antenna, contralateral to the plate, was ablated at the base. After the ablation, the animal was left undisturbed for more than 40 minutes, after which its movements were recorded under the control and wall conditions. For each condition, five trials were performed with an inter-trial interval greater than one minute. In total, 20 trials were performed for each individual. Twenty-four individuals were used.

## Data analysis

Behavioral data provided by the TrackTaro software were processed and analyzed offline, using algorithms customized using Python 3.7.1 (Jupyter Notebook version 5.7.4). First, we classified all recorded data from all trials into “wind-elicited response” or “no response” based on the walking speed, as used in previous studies using the treadmill system (Oe and Ogawa, 2013; Fukutomi et al., 2015; Fukutomi and Ogawa, 2017). The response probability was defined as follows:

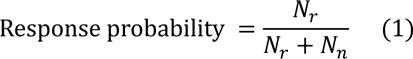

Where, *N*_*r*_ and *N*_*n*_ are the number of trials categorized as a wind-elicited response and a no response obtained for each stimulation condition, respectively. We focused on the initial responses to the air-puff stimulation, which was defined as a continuous walking trot followed by a stationary moment, and we measured four locomotion parameters: walking direction, turn angle, reaction time, and walking distance. Definition and calculation of these parameters are similar to those in our previous study (Oe and Ogawa, 2013; Fukutomi et al., 2015; Fukutomi and Ogawa, 2017). The walking direction and turn angle were defined for the forward direction as 0°, clockwise as plus, and counterclockwise as minus. We arranged the walking direction so that it ranged from 0° to ±180°. To compare the magnitude of the turning movement, unlike the walking direction, the turn angle was not arranged. The trajectory length of the initial response is referred to as the walking distance. The forward distance was defined as the travel distance of the forward movement in the initial response. The reaction time was defined as the interval from the cricket’s receiving the airflow to the start of the initial response. To calculate the reaction time, the travel time of the airflow between the nozzle and the cricket was subtracted from the time delay from the opening of the delivery valve in the picopump to the start of the response (Fukutomi et al., 2015).

### Statistical methods

R programming software (version 3.5.2, R Development Core Team) was used for the statistical analysis. To avoid pseudo-replication, we used the mean value of the data obtained in the trials categorized as wind-elicited response for each individual as the representative value for the statistical tests. Because the walking direction is a circular parameter, we calculated the circular mean angle of the walking direction for each individual. Prior to statistical testing, we checked the distribution of the dataset for the walking distance, turn angle and the reaction time, using the Shapiro-Wilk test. As the data for both parameters in all the experiments were not normally distributed, we used the Wilcoxon rank-sum test if all tested-individuals did not respond or Wilcoxon signed-rank test if all tested-individuals responded, to assess the significance of the stimulation conditions for walking distance, turn angle, and reaction time. The Wilcoxon signed-rank test was also used to assess the significance of the stimulation condition for the response probability. Fisher’s nonparametric test for the common median direction (Fisher, 1993; Pewsey et al., 2013) was used to assess the significance of the stimulation condition for the walking direction. To assess the significance of the stimulation condition for the circular homoscedasticity of the walking direction, we used the Wallraff test, for which the package “circular” was used (Agostinelli and Lund, 2017). All statistical tests for the significant effects among three or more groups were corrected with Bonferroni correction.

## RESULTS

### Effects of antennal mechanosensory inputs on the cercal-mediated escape behavior

A short air-puff elicits running or jumping in crickets, which is considered an escape behavior because they move in the direction opposite to the stimulus (Oe and Ogawa, 2013; Fukutomi et al., 2015; Sato et al., 2019). The cercal sensory system, which is an abdominal mechanosensory organ, mediates wind-elicited escape behavior. First, to test whether the cricket’s antennae also could sense the airflow stimulus and contribute to the escape behavior, we compared the response to the air puff of intact and bilateral antennae-removed crickets. If the antennal mechanosensory inputs were involved in the wind-elicited escape behavior, ablation of the antenna would alter the response to the stimulus from the anterior because the antenna was more sensitive to the frontal stimulus. Here, the air-puff stimulus was applied from eight angles around the cricket, and the stimulus-angle-related directionality in the escape behavior was examined. The antenna-ablated crickets responded to the air-puff stimuli applied from any angle, and their trajectory did not differ from that of intact crickets (Fig. S1A). The antenna ablation had little effect on either the walking direction or the turn angle for any stimulus angle (Fig. S1B–D). There was also no significant difference in angular dispersion around the mean angles of the walking direction between the cut and intact conditions (P > 0.05, Wallraff test). The reaction time was also not affected by antennal ablation (Fig. S1D). In contrast, the turn angle, walking distance, and response probability were affected only for the specific stimulus angle (Fig. S1D). This suggests that antennal ablation would slightly reduce motor activity but had little impact on responsiveness and directional control. In conclusion, our results indicate that the antenna stimulated by the airflow did not contribute to the wind-elicited escape behavior of crickets.

Next, we tested whether the crickets could alter the escape behavior when they actually detected obstacles with their antenna. We also examined how the crickets modulated the escape trajectories depending on the object shape and distance. Fig. 1 shows the responses to the air puff from behind when a cylindrical pole or square plate was positioned at different distances (Fig. 1A). The walking trajectories showed that the crickets tended to walk toward the free side, opposite to the objects, and this movement was most pronounced in the near wall condition (Fig. 1B). The data on the walking direction revealed that it was significantly biased toward the contralateral side of the plate in the near wall condition (Fig. 1C, E). In contrast, the pole had no effect on the walking direction, regardless of its location. The turn angle, which is the angle at which the cricket was rotated in the initial response (Oe and Ogawa, 2013), appeared to increase the rotation toward the contralateral side to the plate positioned at the near position (Fig. 1D). However, there was no significant effect of either the pole or plate on the turn angle (Fig. 1E). The reaction time and walking distance, and response probability, which were non-directional parameters, were not affected by the presence of objects.

We further tested the effects of the pole or plate placed at different distances on the response to airflow applied from the lateral side (Fig. 2A). In the control condition with no object, the cricket moved in the direction opposite to the air-puff stimulus, as shown by gray traces in Fig. 2B, and the walking trajectories were distributed around –90° (Fig. 2C). However, when the objects were placed on the contralateral side of the stimulus, the cricket moved more backward and the distributions of the walking direction were shifted (Fig. 2C). The backward bias in the walking direction was significant when the plate was positioned at the near position (Fig. 2E). The plate located at the far position also tended to bias the walking direction backwards, but this effect was not significant. The pole did not affect the walking direction, regardless of the location. These results suggest that crickets alter their walking direction to avoid hitting the obstacle depending on the obstacle shape and distance to the obstacles. The modulation in the walking direction would solely result from antennal contacts but not from air turbulence by the objects because the effect of the plate that was located at the near position was abolished by bilateral ablation of the antennae so that the crickets could not detect it (Fig. S2). In contrast, neither the pole nor plate had any effect on the turn angle, walking distance, and response probability regardless of their location (Fig. 2C, E). The plate placed at the near position tended to increase the reaction time, but this effect was not significant. To avoid collision with the wall, the cricket might have to delay the onset of escape response. There was a significant difference in the reaction time between control conditions 1 and 2, which were performed without objects before and after the experiments using other conditions with objects (Fig. 2E). This might be due to the after-effects of previous encounters with obstacles. In summary, the results shown in Figs 1 and 2 indicate that the crickets were able to sense the object shape and the distance to it and to modulate escape trajectory to avoid collision with obstacles by using their antennal mechanosensory system.

### Crickets could sense the location of obstacles with their antenna

Next, to test the ability of the antennal system to sense the location and orientation of the obstacles, we examined the effects of the location of the plate on the escape behavior. First, we placed the plate on the lateral side of the cricket at two different positions, anterior or posterior (Fig. 3A). We predicted that crickets would change their movement direction depending on the plate location: they might move backwards when the plate is placed at the anterior position and forward when it is placed at the posterior position. The stimulus was applied from the contralateral side to the plate.

The walking direction was biased backwards by the anterior plate. In contrast to our prediction, the posterior plate had no significant effect and did not enhance forward movement (Fig. 3B, C, E). This bias in the walking direction was coupled with a longer walking distance. It is likely that crickets might have to move longer distances in order to avoid the anterior wall (Fig. 3E). The turn angle, reaction time, and response probability were not affected by the presence of the plate in either positions (Fig. 3D, E). The lack of an effect of the plate positioned posteriorly might be due to the cricket’s failure to detect it. To confirm this possibility, we observed the voluntary movement of the antenna against the plate at the posterior position by using a high-speed camera. There were no significant differences in either the duration or frequency of the antennal contacts between the anterior and posterior positions of the plate (Tables S1, S2). This observation indicated that the cricket could perceive the plate placed at the posterior position. The changes in their escape behavior depending on the wall position suggest that crickets can sense the location of the objects with the antenna.

### Bilateral antennal inputs were necessary to sense the position of frontal obstacle

The results thus far revealed that crickets could change their escape behavior by locating objects even with only one antenna. This was because it was difficult for the crickets to touch the laterally placed plate and pole with their contralateral antenna. If both antennae were used to detect the object, the crickets could perceive the object location more precisely. To examine this possibility, we studied the trajectory of the escape response to the airflow from the rear when the plate was placed in front of the cricket at different positions namely, at the center or to one side (Fig. 4A). In this experiment, the crickets were able to touch the plate at the front center with both antennae, while the plate located to one side could be touched by only the ipsilateral antenna. We predicted that the crickets in response to the airflow from the rear would move forward, biased to the free side and contralateral to the plate, when the plate was placed at one side. As expected, the plate placed on one side significantly biased the walking direction and turn angle to the free side opposite to the wall (Fig. 4B–E).

On the other hand, when the wall was placed in front of the animal and aligned to the center, the trajectory was greatly altered. Although there was no significant difference in the mean walking direction and the turn angle compared to the control condition, both directional parameters became widely distributed and the number of responses decreased around 0° of the walking direction and turn angle (Fig. 4C–E). Furthermore, there was no significant difference in the walking distance between the conditions, but the forward movement was significantly reduced in the center wall condition (Fig. 4E). These results indicate that the center wall might enhance the lateral movement rather than the forward movement. The cricket could alter its escape movement depending on the position of the obstacle to effectively avoid collision with it. In addition, the reaction time significantly increased when the plate was placed centrally in front of the animal (Fig. 4E). It might take a longer time for the cricket to make a decision on the escape direction. The plate placed in the frontal region of the cricket had no effect on the response probability regardless of its position, suggesting that the obstacles on the escape route might not suppress the escape decision.

It was revealed that crickets could modulate their escape behavior depending on the position of the obstacle. This then leads us to the question: did the crickets need to compare the left and right antennal inputs to sense the precise plate position? To answer this, we ablated an antenna contralateral to the plate and examined the modulation of the escape behavior triggered by the stimulus applied from behind when the plate was positioned in the front, ipsilateral to the intact antenna (Fig. 5A). Unilateral ablation of the antenna had no effect on the escape in the absence of walls (control in Fig. 5). However, when the plate was placed contralateral to the ablated side, the biased effects on the walking direction and turn angle observed in intact crickets were abolished (Fig. 5B–E). This result indicates that bilateral antennal inputs were required to sense the position of the frontal obstacle. To determine if the intact crickets could contact the plate with their opposite antenna, we monitored the antenna movement using a high-speed camera. We found that intact crickets rarely touched the plate with the contralateral antenna (Tables S3, S4). These results imply that even when an antenna is not making contact with the object, it might be providing some signals useful for recognizing the object location.

## DISCUSSION

### Object recognition by antennal mechanosensing

In this study, the crickets were tethered on the treadmill so that the experimenter could manipulate the shape, distance, and position of the stimulating objects. The crickets modulated their trajectory depending on the distance to the object, the object shape, and its orientation. The plate at the near position altered the walking direction, but that at the far position did not. In contrast, the pole had no effect regardless of the distance from the animal (Figs 1 and 2). In addition, the effects of the laterally placed plate differed depending on its anterior-posterior location (Fig. 3), and the plate placed in front of the cricket had different effects on the escape response from that placed laterally (Fig. 4). These results indicated that the crickets were able to perceive not only the objects that they came in contact with, but also their spatial information, including distance, location, and orientation.

Cockroaches use the antennal system to guide their locomotion in an environment with obstacles such as barriers and walls (Camhi and Johnson, 1999; Harley et al., 2009; Baba et al., 2010). When cockroaches actively touch an object placed close to their antenna, they exhibit different turning behaviors depending on the horizontal position of the object (Okada and Toh, 2000, 2006). These facts suggest that the antennal system in cockroaches is a mechanosensory system that is able to identify the location and orientation of objects. However, if an object detected by the antennae directly elicits an action that is performed in a certain relationship to the object position, the cockroach can avoid or localize an obstacle. Thus, it is difficult to distinguish whether this orienting behavior is caused by the perception of the entire surrounding space or simply by a reflexive response to the tactile stimulation. In contrast, our present study directly demonstrates the spatial recognition ability of the crickets’ antennal system by examining the escape behavior elicited by airflow stimulus, which was not mediated by the antennae (Fig. S1). Crickets altered their escape behavior even for identical airflow stimulus applied in the same direction, depending on the location and orientation of the object. This suggests that the crickets may be able to perceive the entire arrangement of objects in the surrounding space. The cricket, which was tethered on the treadmill, actively sensed the objects by moving its antennae freely and scanning the surrounding space. This has also been reported in cockroaches (Okada and Toh, 2006). Active sensing by a cricket’s antennal system provides advanced spatial information to perceive the surrounding space, which will allow the crickets to modulate their behavior flexibly.

### Context dependent modulation of escape behavior

The presence of an object detectable by the antennae altered the direction in which the cricket moved in response to the airflow. This wind-elicited movement is considered to be one of the escape behaviors in which the cricket perceives the airflow as a cue for the approach of a predator (Tauber and Camhi, 1995; Casas and Dangles, 2010; Dupuy et al., 2011). The direction of the wind-elicited escape in crickets is precisely controlled depending on the stimulus angle, similar to the rapid escape in flies and cockroaches (Domenici et al., 2008; Card, 2012). In the absence of an object, as in the control condition, the crickets fled in the opposite direction from which the air-puff stimulus came (Fig. S1; Oe and Ogawa, 2013). Consistent with previous studies, the directions of movement in the initial response to the airflow from either the rear or lateral side that was used in this study were also distributed around the front or contralateral direction to the stimulus (Fukutomi et al., 2015; Sato et al., 2019). However, when a plate was placed on the antero-lateral side of the cricket, direction that the cricket moved in was shifted to the opposite side of the plate for stimuli that were applied from the rear (Fig. 1), and the movement was shifted backwards for lateral stimuli, that were applied from the opposite side of the plate (Fig. 2). These shifts in the walking direction are thought to indicate avoidance of collisions with objects perceived as walls. Cockroaches walking near a wall maintain a constant distance while keeping their antenna in contact with the wall (Camhi and Johnson, 1999), and surgically shortening the cockroach’s antenna increases the collision rate to the wall (Baba et al., 2010). The results from the present study show that the crickets flexibly alter their escape trajectory to avoid collisions, depending on the angle of airflow (stimulus), even if the same obstacles are placed in an identical position. For example, when the plate was placed at the near position on the lateral side of the crickets, their forward movement triggered by the stimulus applied from behind was biased toward the opposite side of the plate, while the lateral movement induced by the stimulus from the opposite side of the plate was altered toward the back (Fig. 3, 4). This suggests that crickets could modulate their behavior depending on the spatial relationship between the stimulus directly triggering the behavior and the environment, that is, the spatial context. It has been reported that descending signals from the cricket brain are necessary to regulate the escape direction (Oe and Ogawa, 2013). Some descending neurons sensitive to artificially caused antennal movement have been identified in the cricket brain (Gebhardt and Honegger, 2001). The mechanosensory information presented by the insect’s antennal system is possibly processed by the brain and used to control their movement for successful escape via descending neurons.

The changes in escape trajectory depended on the location and orientation of the objects. The crickets moved toward a free space in the absence of obstacles (Figs 3, 4). This fact suggests the ability of crickets to perceive the arrangement of objects in the surrounding space. Interestingly, the escape response to the airflow from the rear was delayed and the forward movement was suppressed when the plate was positioned at the center in front of the cricket (Fig. 4E). The crickets might have hesitated to run forward in order to avoid a collision with the wall in the front and delay their decision making before starting their escape movement. In contrast, the plate positioned to one side in front of the animal biased the escape toward the wall-free side but did not affect the reaction time. This implies that the response delay and the reduction in the moving distance are not simply due to the reactive inhibitory effects of the mechanosensory inputs from the antennae. It has been reported that high-frequency sounds hinting at the presence of predators reduce the response probability of wind-elicited escape (Fukutomi and Ogawa, 2017). Our results imply that crickets can flexibly change their escape behavior depending not only on acoustics but also on the spatial contexts sensed by the antennal system in the surrounding space. The crickets integrate the sensory inputs of multiple modalities to perceive the surrounding context and make decisions for an appropriate and successful escape.

### Crickets sensed “absence” of objects in surrounding space by their antennal system

Comparing the inputs from the left and right antennae is useful for obtaining a more accurate picture of the surrounding environment. For example, in navigation to localize an odor source, insects sample chemicals using a pair of antennae as chemical sensors and compared the sensory inputs to orient (Eiras and Jepson, 1994; Willis, 2008; Takasaki et al., 2012). The bilateral comparison of antennal inputs has also been reported for mechanosensory cues. Crayfish have been reported to compare tactile inputs from both antennae to determine the turning direction (McMahon et al., 2005). Our results indicated that crickets compared tactile information between the left and right antennae to move toward the obstacle-free space, because unilateral ablation of the antenna contralateral to the frontal plate diminished the effects on the direction of movement and turn angle (Fig. 5E). This meant that the antenna on the object-free side provided information of “no object” even if it did not contact anything. Since this information was no longer provided by the ablated antenna, it would not be possible for the crickets to accurately localize the object detected by the intact antenna. The perception of the absence of objects is one of the important functions of active sensing, for which animals repeatedly scan the environment by moving their sensory organ voluntarily (Bermejo et al., 2005; Nelson and MacIver, 2006). Our results suggest the possibility that insects can recognize the absence of objects by using active sensing with their antennal mechanosensory system.

## Competing Interests

The authors declare no competing interests.

## Author contributions

N.O.I., H.S., and H.O. designed research. N.O.I. performed experiments. N.O.I and H.S analyzed all data. All authors discussed the data interpretation and statistical methods. N.O.I., H.S., and H.O. wrote the manuscript.

## Funding

This work was supported by funding to H.O. from JSPS and MEXT KAKENHI grants 16H06544.

## Data Availability

The data that support the findings of this study are available from the corresponding author, H.O., upon reasonable request.

## Notes

### Competing Interest Statement

The authors have declared no competing interest.

